# Ozone mediates tumor-selective cell death caused by air plasma-activated medium independently of NOx

**DOI:** 10.1101/2023.03.17.533239

**Authors:** Manami Suzuki-Karasaki, Yushi Ochiai, Shizuka Innami, Hiroshi Okajima, Miki Suzuki-Karasaki, Hideki Nakayama, Yoshihiro Suzuki-Karasaki

## Abstract

Cold atmospheric plasma and plasma-treated liquids (PTLs) are emerging promising tools for tumor-targeted cancer treatment, as they preferentially injure tumor cells more than non-malignant cells. Oxidative stress is critical to the antitumor effect, but the oxidant mediating the effect is debatable. Previously, we reported that air plasma-activated medium (APAM) has tumor-selective cytotoxicity in vitro and in vivo. Moreover, an unusual mitochondrial positioning named monopolar perinuclear mitochondrial clustering (MPMC) and nuclear damage proceeds to cell death. We noticed that air plasma generation was accompanied by ozone (O_3_) formation, leading to suppose the possible role of O_3_ in the effect of APAM. In this study, we produced an O_3_-dissolved medium (ODM) and comparatively analyzed its biological effect with APAM. Both agents had comparable amounts of dissolved O_3_ (dO_3_), while APAM, but not ODM, contained nitrite and nitrate. Like APAM, ODM could induce apoptosis, nonapoptotic cell death, tubulin remodeling, MPMC, and nuclear shrinkage. Catalase mitigated all these events. The increases in various intracellular and mitochondrial reactive oxygen species (ROS) and lipid peroxides proceeded to cell death, and catalase also prevented them. Conversely, suppressing cellular H_2_O_2_ removal systems augmented mitochondrial ROS production and cell death. In contrast, like APAM, ODM minimally increased ROS production and MPMC in non-malignant cells. These results indicate that dO_3_ is a critical mediator of the actions of APAM, including tumor-selective induction of MPMC and cell death. Our findings suggest ODM could be a more chemically-defined alternative to PTLs in cancer treatment.

## 1. Introduction

Like other intractable cancers, once developed, oral cancers (OCs) are highly refractory, recurrent, metastatic, resistant to anticancer agents and irradiation, and hard to remove by surgical resection [1,2]. Thus, improving prognosis through standard therapies remains challenging, and innovative treatment is urgently required. Cell death induction is one of the powerful strategies to eliminate cancers. Apoptosis is the representative mode of cancer cell death targeted by conventional therapies, including most anticancer agents and radiation. However, intractable cancers, including OCs, have congenital and acquired resistance. It is widely accepted that besides apoptosis, various nonapoptotic cell death forms contribute to cancer cell death [3]. Consequently, drugs and tools targeting other forms of cell death could serve as successful venues for OC treatment.

Cold atmospheric plasmas (CAPs) are partially ionized gasses containing ions, electrons, and free radicals and have tumor-selective toxicity. Flushing CAPs to various solutions, such as culture media, buffers, and infusion fluids, result in plasma-treated liquids (PTLs), which also preferentially injure tumor cells while damaging non-malignant cells minimally [4–6]. CAPs and PALs also reduced tumor growth in animal models with minimal adverse events. Thus, CAP-based treatments have attracted much attention in cancer treatment. However, PTLs have different chemical and biological properties, depending on the physical nature of the plasmas generated, the solutions used, and various experimental parameters. In addition, the biological outcomes may also vary significantly depending on the target cell systems. As a result, their actions, including targeted cell death, are highly complicated; While most of them induce apoptosis, some of them can trigger nonapoptotic cell death forms, including autophagy [7.8], necroptosis [6], and ferroptosis [9] in various cancer cell models. Although numerous reports have shown the vital role of reactive oxygen and nitrogen species (ROS/RNS) in mediating the antitumor effect, the detailed mechanisms remain obscure. Moreover, PTLs have complicated components, and complete chemical identification of the active substances is challenging. Indeed, many kinds of chemical species, atomic oxygen (O), singlet oxygen (^1^O_2_), superoxide (O_2_^·-^), hydroxyl radicals (^·^OH), hydrogen peroxide (H_2_O_2_), nitric oxide (NO^·^), and nitrogen oxide anions (NOx^-^), such as nitric dioxide (NO_2_^·^), nitrite (NO_2_^-^), and nitrate (NO_3_)^-^ have been proposed to participate in the effect. More complicatedly, the exact contents of these species vary considerably depending on the CAP source, settings, and ambient conditions. In addition, these oxidants can function cooperatively. Several studies have shown the synergistic effect of H_2_O_2_ and NO_2_^-^ in cell death and tumor selectivity [10, 11]. Consequently, the primary mediator(s) and their molecular targets in the antitumor properties are poorly understood. Such chemical ambiguity heavily frustrates distinct descriptions of the mechanism of action and clinical application.

O_3_ is the oxygen allotrope consisting of three oxygen atoms. It is an acrid gas at ordinary temperature and pressure. O_3_ is an unstable molecule with unique physicochemical and biological properties due to its resonance structures. It has the second-highest oxidizing power behind fluorine. O_3_ directly injures the cell membrane of bacteria through the oxidation of phospholipids and lipoproteins and has potent antibacterial activity. O_3_ also inactivates viruses, fungi, yeast, and protozoa. Therefore, it has been widely used for sterilization and disinfection. O_3_ has also been utilized in treating various diseases for over a century [12]. Moreover, numerous in vitro and in vivo studies and a few clinical trials [13–15] have shown the anticancer effects of O_3_. O_3_ has a direct antitumor effect in some cancers while having indirect effects, such as immunomodulation, synergistic or adjuvant effects with various anticancer drugs and radiation (cisplatin, 5-fluorouracil, etoposide, and gemcitabine) [16–18]. O_3_ is administered to animals in various ways, including topical gas, rectal or intraperitoneal insufflation, and intravenous or intratumoral injection. O_3_ has a substantial solubility in water (14 mmol/L at 20°C) and has also been used with nanotechnology as ozonated or ozonized water. However, it must be aware that pure O_3_ production by conventional methods is challenging. The most widely used method to generate O_3_ is the silent discharge in the air. However, this method results in the excitation of molecular nitrogen (N_2_) as well as molecular oxygen (O_2_), leading to the production of various nitrogen oxides (NOx). Accordingly, ozonated/ozonized fluids could contain a mixture of dO_3_ and multiple bystander nitrogen oxide anions (NOx^-^). NOx, such as nitric oxide (NO^·^) and nitric dioxide (NO_2_^·^), can produce O_3_ and more complicated, harmful oxidants in the presence of light. In addition, NO^·^ has been recognized as a primary modulator of cell death and a potential target for anticancer therapy [19, 20]. As a result, the co-existence of bystander NOx could confuse the precise evaluation of the biological activity of dO_3_ in these ozonated fluids. Therefore, eliminating NOx is essential to define the effect of dO_3_ itself.

Mitochondria are highly plastic, dynamic, and heterogenous organelles in connection with these diverse functions. Their size, shape (macroscopic shape), and location vary in different tissues, cells, and experimental conditions. An emerging view is that mitochondrial shape and location changes are not passive events. Instead, they are active events coupled with cellular functions, cell death, and survival [21–23]. Mitochondria can distribute broadly throughout the cytoplasm (Pan-cytoplasmic), subplasmalemmal, or perinuclear sites. Pan-cytoplasmic distribution governs Ca^2+^ transport from the endoplasmic reticulum (ER) via tethering with ER [24]. The subplasmalemmal distribution controls Ca^2+^ channel activity by regulating mitochondrial function as a Ca^2+^ reservoir [25]. The mitochondrial location around the perinuclear regions is called perinuclear mitochondrial clustering (PNMC). This response is induced by stresses, such as hypoxia and heat shock, and is associated with mitochondrial fragmentation [26, 27]. This type of positioning is thought to be adaptive to stresses and cytoprotective. Microtubule polymerization inhibitors, such as Nocodazole (NC) and Cholchicine, and the expression of KinAΔN355 mutant of Kinesin prevent it, indicating that microtubule- and Dynein and Kinesin-dependent. Mitochondria may be transported along the microtubule track. PNMC participates in cancer adaptation to hypoxia through mitochondrial reactive oxygen species (ROS) production and Hypoxia-inducible factor-1α, the master transcription factor of hypoxic signaling [26]. Recently, we reported another type of mitochondrial distribution associated with cancer cell death [28]. In this case, mitochondria become fragmented and gather one side of the perinuclear sites. Drastic changes in the shape and location of tubulin accompany this phenomenon. Like mitochondria, tubulin concentrates one side of the nuclei. NC and antioxidants prevent these changes, suggesting the involvement of similar mitochondrial transport mechanisms in PNMC and ROS. We named this unique mitochondrial positioning monopolar perinuclear mitochondrial clustering (MPMC) to distinguish it from other known mitochondrial locations.

We noticed that air plasma generation was accompanied by O_3_ production. This notion led us to assume the possible role of the oxidant in the effect of APAM. To examine this hypothesis, we developed a new dO_3_-generating system to produce O_3_ without exciting N_2_ and dissolved it in a culture medium by bubbling. The resulting O_3_-dissolved medium (ODM) can keep dO_3_ for a month at -80°C. In this study, we comparatively analyzed the biological effects between APAM and ODM. Results showed that without NOx^-^, dO_3_ could mimic various effects of APAM, including tumor-selective induction of MPMC and oxidative cell death.

## 2. Materials and methods

### 2.1. Materials

Unless otherwise specified, all chemicals were parched from Sigma-Aldrich (St. Louis, MO, USA). The pan-caspase inhibitor Z-VAD-FMK was obtained from Merck Millipore (Darmstadt, Germany). All insoluble reagent was dissolved in dimethyl sulfoxide (DMSO), and the stock solution diluted in 10% Fetal Bovine Serum(FBS) contained Dulbecco’s Modified Eagle Medium (DMEM; final DMSO concentration,<0.1%) before use.

### 2.2. Cell culture

The human OC cell line HOC-313 was kindly provided by the Department of Oral and Maxillofacial Surgery, Graduate School of Medical Science, Kanazawa University (Kanazawa, Japan). Another human OC cell line SAS and glioblastoma (GBM) cell line U251MG were obtained by the Japanese Collection of Research Bioresource (JCRB) Cell Bank of the National Institute of Biomedical Innovation, Health, and Nutrition (Osaka, Japan). Human fetal osteoblast hFOB was a kind gift from Dr. T. Ando (Yamanashi University). Human dermal fibroblasts (HDFs) were obtained from Cell Applications (San Diego, CA, USA). The cells were maintained in 10% FBS (Serena Europe Gmbh, Brandenburg, Germany) containing DMEM (Merck, NM, USA) supplemented with 100 U/mL penicillin and 100 μg/mL streptomycin (FBS/DMEM) at 37 °C in a 5% CO_2_ incubator.

### 2.3. O_3_ and ODM preparations

O_3_ was generated in two different ways. Method A produced O_3_ by irradiating an excimer light (185 nm) into the air using a UV-C lamp (Fig. 1A). This method allows specific excitation of O_2_ but not molecular nitrogen (N_2_), the cause of NOx production. Method B made it by the silent discharge of highly pure (99.9%) O_2_ (Taiyo Nippon Sanso JFP Corporation, Kanagawa, Japan) using the O_3_ generator (Communication&Control Systems Company, Tokyo Keiki Incorporation, Tokyo, Japan) equipped with a dielectric barrier discharge probe illustrated in Fig. 1B. This closed system does not allow air contamination and the resulting excitation of N_2_. The generated O_3_ was introduced into phenol red-free DMEM (Fuji Film Wako Chemicals. Osaka, Japan) by bubbling, resulting in an O_3_-dissolved medium (ODM). The typical experiment conditions are a 10–15 KV voltage and a gas flow rate of 0.4 L/min. ODM was made by O_3_ bubbling at the ratio of 1 min bubbling/1 mL ODM. The original ODM was diluted to a final concentration of 6.3–50% with phenol red-free DMEM (biochemical experiments) or FBS/DMEM (for cell experiments).

**Figure 1.**
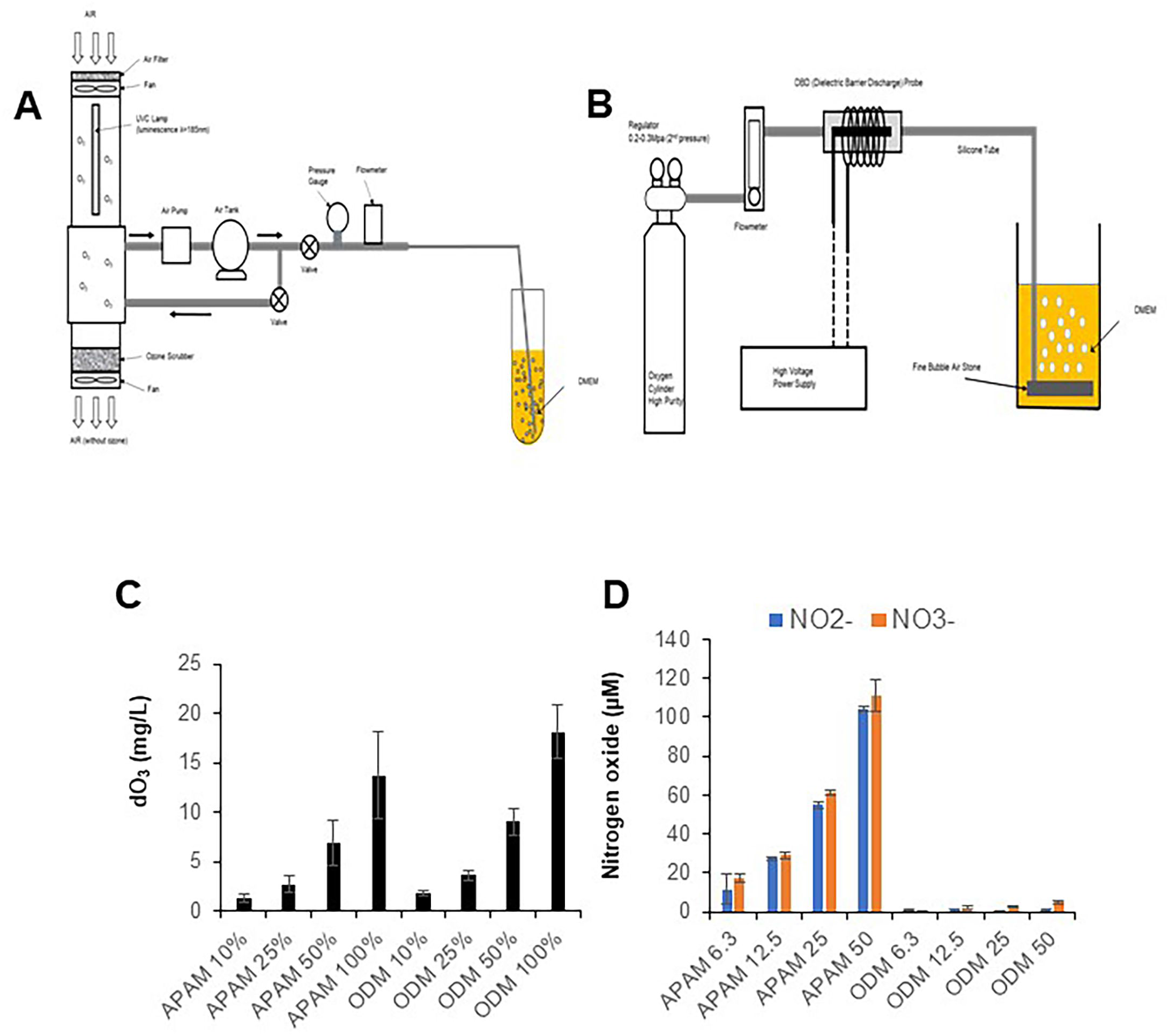
ODM has dO_3_ but not NO_2_^-^/NO_3_^-^. (A, B) Schematic O_3_-generator and bubbling system diagrams. In method A, O_3_ was produced by irradiating an excimer light (185 nm) into the air using a UV-C lamp (A). In method B, O_3_ was generated by exciting high-purity (99.9%) molecular oxygen (O_2_) with dielectric barrier discharge (B). The resulting O_3_ was then introduced into a DMEM by bubbling. (C) Quantitation of dissolved O_3_ (dO_3_) in ODM and APAM. The amount of dO_3_ in ODM and APAM (12.5−50% solution) was measured as described in Materials. Data are the mean ± SD (n=4). (D) The concentrations of NO_2_^-^ and NO_3_^-^ (C) in ODM (6.3–50% solution) were measured using the Griess method. APAM was used as a positive control. NO_3_^-^ concentration was calculated following the formula; [NO_3_^-^] = [(NO_2_^-^ +NO_3_^-^)] − [NO_2_^-^]. Data are the mean ± SD (n=4).

### 2.4. APAM preparation

Air plasma and APAM were prepared as reported previously [28]. Briefly, the plasma was generated from the ambient air using a Piezobrush™ PZ2 model plasma jet (relyon, Germany) equipped with a piezo element. The APAM (1 mL) was made by flushing the plasma to phenol red-free DMEM for 1 min at a distance of 20 mm. The original APAM was diluted to a final concentration of 6.3–50% with the medium (biochemical experiments) or FBS/DMEM (for cell experiments).

### 2.5. Quantitation of oxidants

dO_3_ concentration was measured by a polarographic O_3_ meter (DOZ-1000PE, Custom Corporation, Tokyo, Japan). The concentration was also measured by a Digital Pack test (DPM2-O_3_, range 0.25∼5 mg/L, Kyoritsu Chemical-Check Lab. Corp., Kanagawa, Japan). The concentrations of NO_2_^-^ and NO_3_^-^ were measured using a Kit (NO_2_ / NO_3_ Assay Kit-CII(Colorimetric) Griess Reagent Kit, Dojindo, Kumamoto, Japan) according to the manufacturer’s protocols. The concentrations of NO_2_^-^ and NO_3_^-^ were calculated using a standard curve made using the authentic samples from the kit.

### 2.6. Cell viability assay

Cell viability was measured by the WST-8 assay using a Cell Count Reagent SF (Nacalai Tesque, Inc, Kyoto, Japan). Cells in FBS/DMEM were plated at 4 × 10^3^ cells/well in 96-well plates (Corning Incorporated, NY, USA). They were incubated at 37 °C overnight, added with agents, and incubated for 72h. In the resistance assay, agents were added after incubating cells for 72 h. This prolonged preincubation allows cells to reach a higher density and become less sensitive to agents. After treatment, cells were added 10 μL Cell Count Reagent SF and further incubated for 2h. Absorbances at 450 nm were measured by a Nivo 3F Multimode Plate Reader (PerkinElmer Japan Company, Ltd., Yokohama, Japan).

### 2.7. Determination of cell death

Cell death was evacuated by fluorescence microscopy as previously described [29] with minor modifications. Briefly, cells in FBS/DMEM were seeded at 1.5 × 10^4^ cells/well in a 35 mm Poly-Lysine-coated glass bottom dish (D11531H, Matsunami Glass Ind. Corp., Osaka, Japan) and treated with agents at 37 °C overnight in a 5% CO_2_ incubator. After removing the medium by aspiration, the cells were stained with 4 μM each of Calcein-AM and Ethidium bromide homodimer-1 (EthD-1) for 30min to label live and dead cells, respectively, using a kit (LIVE/DEAD Viability/Cytotoxicity kit) Cells were washed and immersed in FluoroBrite™ DMEM. Images were obtained using BZ-X 710 digital Biological Microscopy (Keyence, Osaka, Japan).

### 2.8. Imaging of mitochondria and tubulin

The mitochondrial morphology and positioning and tubulin were analyzed as previously described [28]. Briefly, cells FBS/DMEM were seeded at a 5 × 10^4^ /mL density in a 35 mm Poly-Lysine-coated glass bottom dish, as described above, and treated with agents at 37 °C for 2 or 18 h in a 5% CO_2_ incubator. After removing the medium, the cells were stained with 20 nM MitoTracker™ Red CMXRos (MTR) or MitoTracker™ Green FM (MTG, ThermoFisher). The nuclei were counterstained with 1 mg/mL of Hoechst33342 for 1 h at 37 °C in a 5% CO_2_ incubator. Tubulin was stained with Tubulin Trackerr™ Green (TTG, ThermoFisher). After washing with FluoroBrite™, the cells were immersed in FluoroBrite™. Images were obtained using a BZ X-710 Digital Biological Microscope (Keyence, Osaka, Japan) equipped with a 100 ×, 1.40 n.a. UPlanSApo Super-Apochromat, coverslip-corrected oil objective (Olympus, Tokyo, Japan) and analyzed using BZ-H3A application software (Keyence). For each experimental group, the mitochondria in 2 or 3 different pictures were counted for three different distribution patterns, pan-cytoplasmic (Type I), PNMC (Type II), and MPMC (Type III).

### 2.9. Measurement of intracellular ROS generation

As described above, cells were seeded in a 35 mm Poly-Lysine-coated glass bottom dish and treated with agents at 37 °C for 2 h in a 5% CO_2_ incubator. After removing the medium, the cells were stained with 1 μM OxiORANGE™ (OXO), HYDROP™ (Hydrop, Goryo Chemicals, Sapporo, Japan), or 5 μM MitoSOX™ Red (MitoSOX, ThermoFisher) to detect ^·^OH, H_2_O_2_, and O_2_^·-^, respectively. Images were obtained using BZ-X 710 digital Biological Microscopy fluorescence microscopy and analyzed using BZ-H3A application software as described above.

### 2.10. Statistical analysis

Data are presented as mean ± standard deviation (SD) and analyzed by a one-way analysis of variance followed by Tuckey’s post hoc test using statistical software with Excel 2019 for windows (SSRI, Tokyo, Japan). P<0.05 was considered statistically significant.

## 3. Results

### 3.1. ODM and APAM have dO_3_

Fig. 1A and B show the schematic diagrams of the generating systems for O_3_ and dO_3_. O_3_ was generated from the air using UV-C excimer light irradiation (Fig. 1A) or from highly pure O_2_ using dielectric barrier discharge (Fig. 1B). The O_3_ was introduced into a culture medium by bubbling, resulting in ODM. Next, we quantified the oxidants in ODM made by method B. As expected, the ODM contained substantial amounts of dO_3_ (Fig. 1C). The quantity was increased over introducing time for at least 30 min. APAM had comparable levels of dO_3_. On the other hand, APAM contained about 100 μM of NO_2_^-^ and NO_3_^-^ while they were under detection limits in ODM (Fig. 1D). ODM made by method A also had dO_3_ but not NO_2_^-^ and NO_3_^-^ (not shown).

### 3.2. ODM can kill different tumor cells

Next, we examined whether ODM affected tumor cell growth. Cells were treated with varying concentrations (12.5– 50% solution) of ODM for 72 h and analyzed for cell growth by WST-8 assay. The treatment dose-dependently reduced the viability of OC cell lines, such as SAS and HOC-313 (Fig. 2A, B). ODM had a similar effect in several OS and GBM cell lines, including HOS (Fig. 2C, D). Cellular sensitivity to ODM varied considerably depending on the cellular growing status. In sensitive cells, ODM (≤12.5%) was enough to reduce their growth potently (≥50% reduction), while in insensitive cells, at higher concentrations (≥25%), it had a moderate effect (<50% reduction). Robust cell and nuclear morphological changes accompanied the growth suppression. Most untreated cells were adherent spindle cells. Treatment with ODM (50%) for 2 h resulted in blebbed, less adhesive round cells, most of which maintained membrane integrity (Fig. 2E, F). At the same time, smooth surface round nuclei became smaller and deformed. These results show that ODM can injure different cancer cell types. We further analyzed the effect of ODM using OC cells as a model.

**Figure 2.**
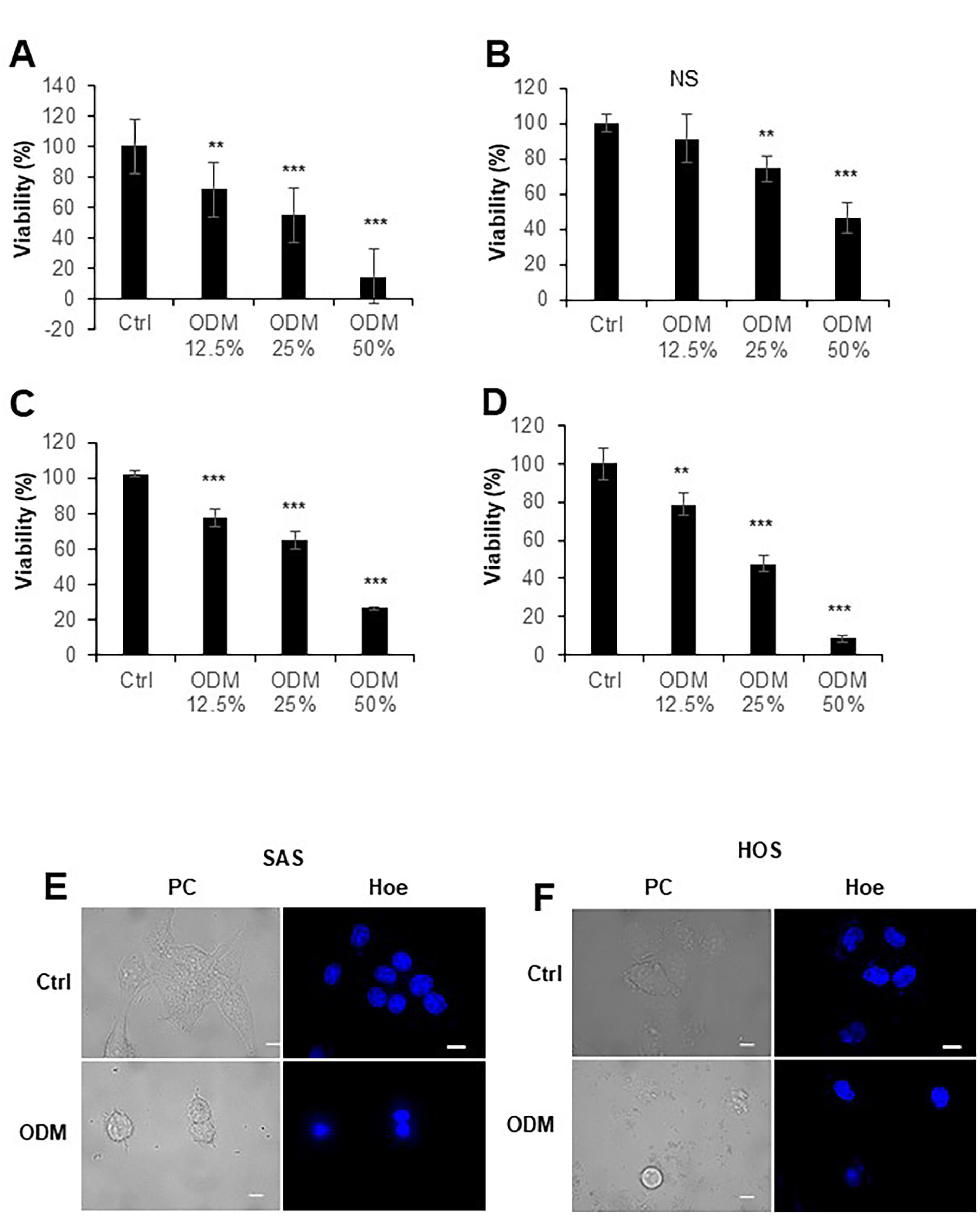
ODM can injure different cancer cell lines. (A−D) SAS (A), HOC-313 (B), HOS (C), and U251MG (D) cells in FBS/DMEM were plated at 4 × 10^3^ cells/well and incubated overnight. The cells were treated with ODM (12.5–50% solution) for 72 h and analyzed for growth using a WST-8 cell growth assay. Data are the mean ± SD (n =8). Data were analyzed by one-way analysis of variance followed by Tukey’s post hoc test. \*\**P* < 0.01; \*\*\**P* < 0.001; NS, not significant vs. control treated with vehicle. (E, F) SAS (E) and HOS (F) cells (1.5×10^4^ cells/well) cells/well were plated in a 35 mm glass bottom dish and incubated overnight, and then treated with ODM (50%) for 2 h. After removing the medium by aspiration, the cells were stained with Hoechst33342 (Hoe). Images were obtained from BZ-X 710 Digital Biological Microscope equipped with a 100x objective and analyzed using BZ-H3A application software. PC, phase contrast. Bar = 10 μm.

### 3.3. ODM primarily induces oxidative cell death in an H_2_O_2_-dependent manner

APAM can induce apoptosis and nonapoptotic cell death in OC cells depending on the cell line and concentration [28]. Therefore, we examined whether ODM had similar effects. When ODM reduced cell viability potently, the broad-spectrum caspase inhibitor Z-VAD-FMK failed to suppress the effect in SAS cells (Fig. 3A). The necroptosis inhibitor Nec-1 did not affect the effect either. We obtained similar results in HOC-313 cells (Fig. 3B). In contrast, Z-VAD-FMK and Nec-1 protected the cells from TRAIL cytotoxicity. The ferroptosis inhibitor Ferrostatin-1 (Fer-1) also failed to affect ODM cytotoxicity (Fig. 3A, B) while preventing Erastin cytotoxicity. On the other hand, when the effect of ODM was moderate, Z-VAD-FMK significantly suppressed it (Fig. 3C), indicating that apoptosis plays a role in specific cellular conditions. Regardless of the degree of the effect, the H_2_O_2_-degrading enzyme catalase protected both cells from ODM, while MnTBaP, the scavenger of O_2_^·-,^ did not (Fig. 3C–E). Catalase also prevented APAM cytotoxicity (Fig. 3F). These results show that similar to APAM, ODM induces oxidative cell death in an H_2_O_2_-dependent manner.

**Figure 3.**
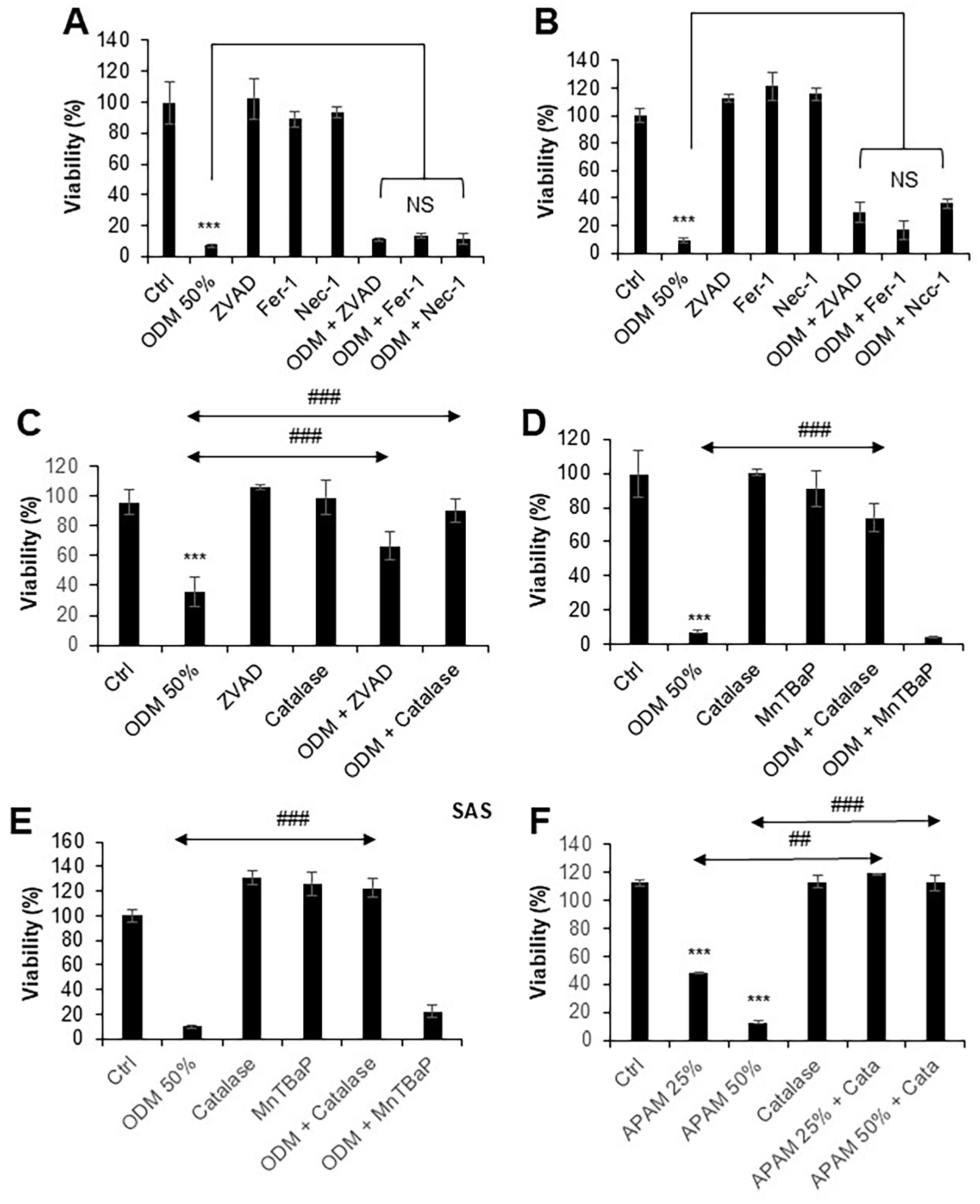
ODM induces oxidative cell death in an H_2_O_2_-dependent manner. (A, B) Effect of cell death inhibitors on ODM cytotoxicity. SAS (A) and HOC-313 (B) cells were processed as described in the legend of Figure 2. The cells were incubated with 10 μM Z-VAD-FMK, Ferrostain-1 (Fer-1), or 30 μM Necrostatin-1 (Nec-1) for 1 h and then treated with ODM (50%) for 72 h. The cell growth was measured as described in the legend of Figure 2. (C) HOC-313 cells were incubated with 10 μM Z-VAD-FMK or 10 U/ml catalase for 1 h and then treated with ODM (50%) for 72 h. The cell growth was measured as described in the legend of Figure 2. Data are the mean ± SD (n =8). Data were analyzed by one-way analysis of variance followed by Tukey’s post hoc test. \*\*\**P* < 0.001 vs. control treated with vehicle. ### *P* < 0.001. (D, E) SAS (D) and HOC-313 (E) cells were incubated with 10 U/ml catalase or 30 μM MnTBaP for 1 h and then treated with ODM (50%) for 72 h. (F) SAS cells were incubated with 10 U/ml catalase for 1 h and then treated with APAM (25, 50%) for 72 h. The cell growth was measured as described in the legend of Figure 2. Data are the mean ± SD (n = 8). Data were analyzed by one-way analysis of variance followed by Tukey’s post hoc test. \*\*\**P* < 0.001 vs. control treated with vehicle. ### *P* < 0.001.

### 3.4. ODM increases mitochondrial oxidative stress in an H_2_O_2_-dependent manner

APAM increases intracellular ROS, including those within mitochondria in OC cells [28]. Therefore, we determined whether ODM also could affect cellular ROS levels. First, we analyzed the intracellular O_2_^·-^ using the oxidant-specific probe MitoSOX Red (MitoSOX) in live cells. O_2_^·-^ was significantly increased in ODM-treated cells compared with untreated cells (Fig. 4A). The increase was observed before significant cellular and nuclear morphological changes. Moreover, catalase abolished the effect, indicating it is H_2_O_2_-dependent. Treatment with the irreversible catalase inhibitor 3-AT increased O_2_^·-^ (Fig. 4A), supporting the role of H_2_O_2_. Figure 4B shows the quantitative analysis of the increase and supports the view. In addition, the treatment increased ^·^OH, detected by OXO, and catalase suppressed the effect, too (Fig. 4C). As MitoSOX and OXO can detect O_2_^·-^ and ^·^OH, respectively, within mitochondria, the above results suggested the increases in mitochondrial ROS (mROS). To further test this view, we analyzed the cellular location of ^·^OH. In untreated cells, the oxidant has broadly distributed the cytoplasm. After ODM treatment, the oxidant was seen at one side of the perinuclear sites. As a result, it colocalized with mitochondria, and catalase abolished the changes (Fig. 4D). These results show that ODM increases mitochondrial oxidative stress in an H_2_O_2_-dependent manner.

**Figure 4.**
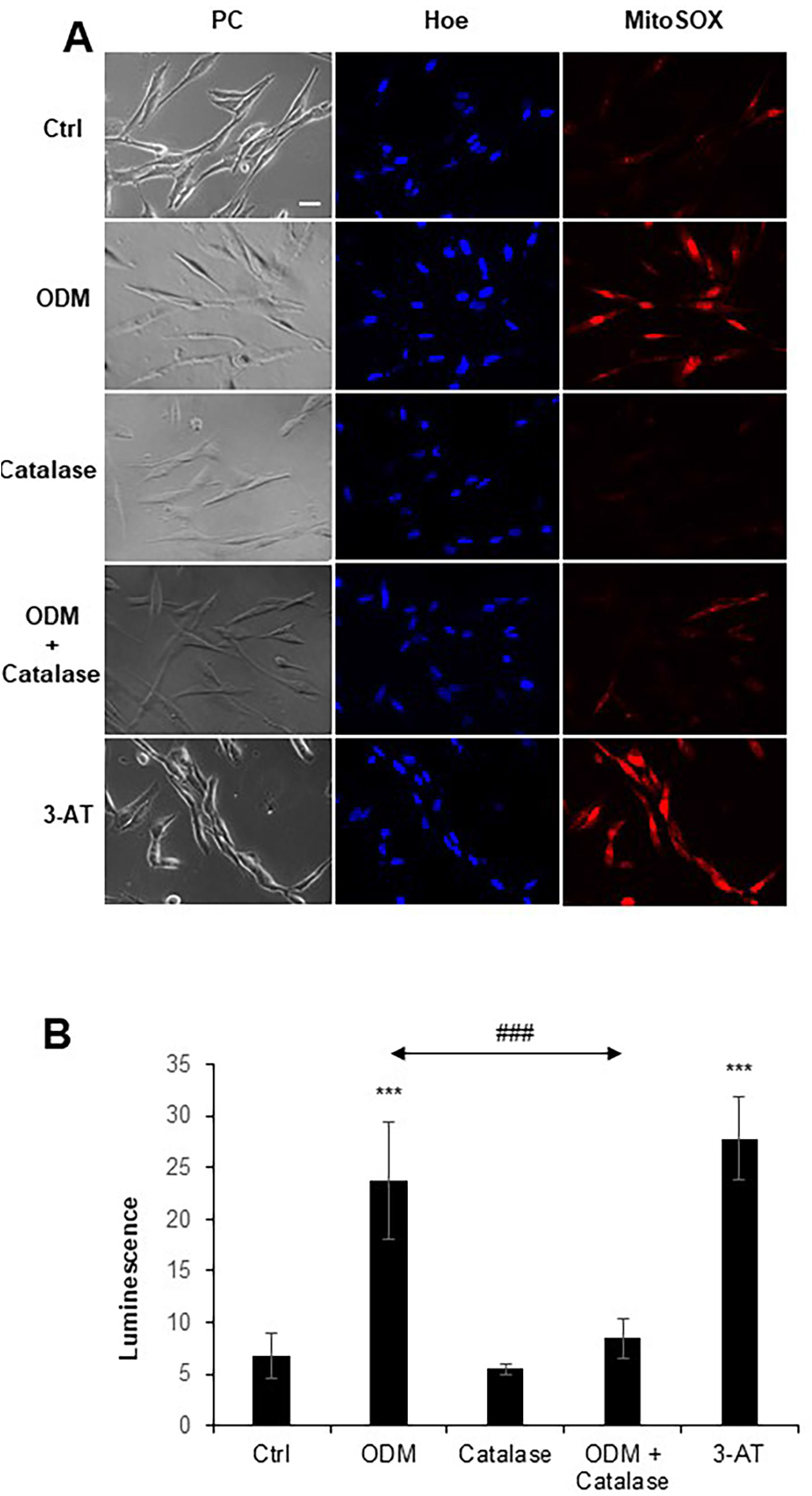
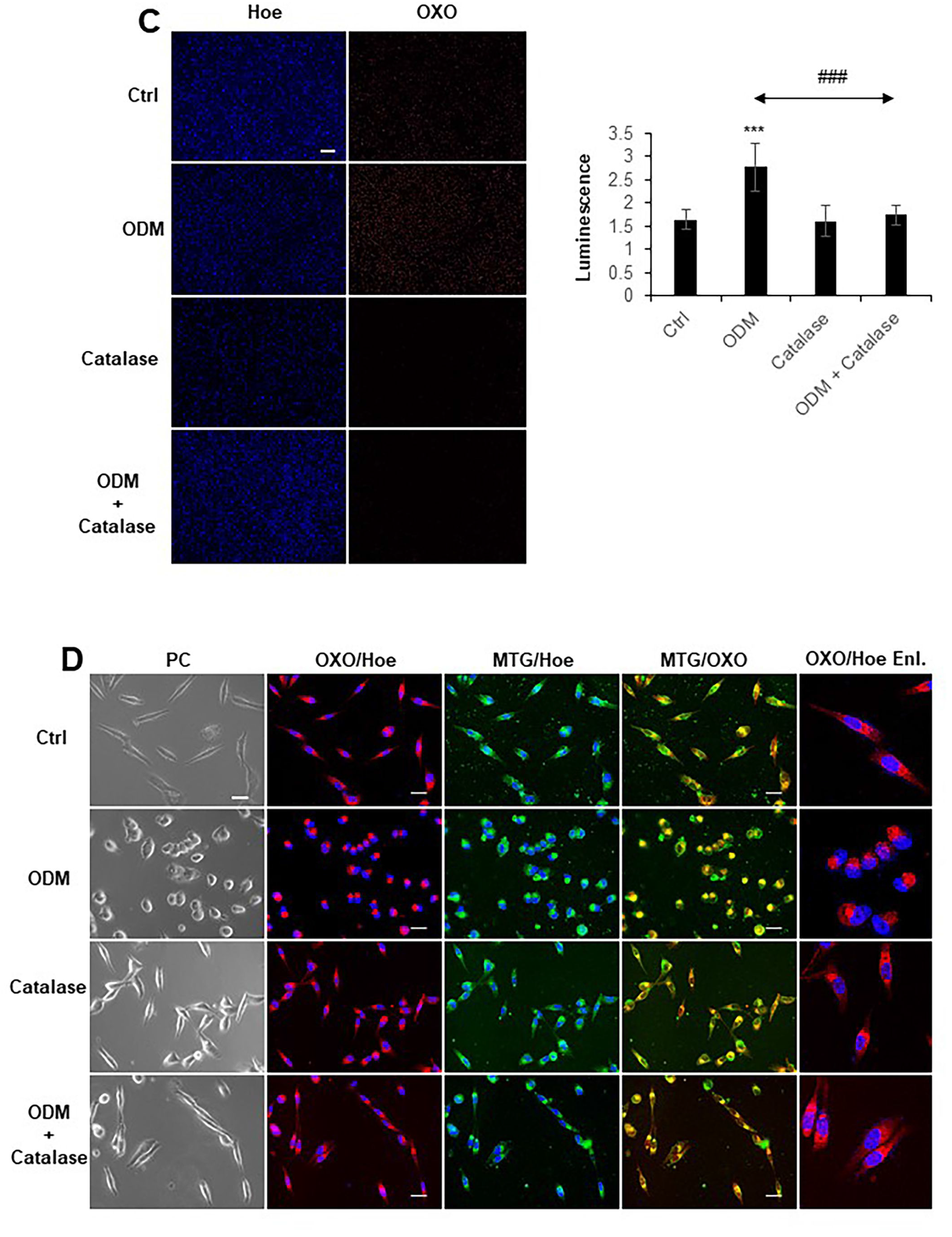
ODM increases different intracellular ROS in an H_2_O_2_-dependent manner. (A) HOC-313 cells were preincubated with 3-amino-1,2,4-triazole (3-AT, 1 mM) and catalase for 1h and then treated with ODM (50%) for 2h. Mitochondrial reactive oxygen species (ROS) and nuclei were stained using specific probes MitoSOX and Hoechst33342, respectively. All images were obtained from BZ-X 710 Digital Biological Microscope with a uniform exposure time (1/2.5S). (B) The luminescence of all color images was measured using a public-domain Java NIH ImageJ program. Data are the mean ± SD (n=9). Data were analyzed by own-way analysis of variance followed by Tuckey’s post hoc test. ###*P* <0.001. (C) The cells were pretreated with catalase for 1h and then incubated with ODM for 2h. Intracellular hydroxyl radicals were stained with OXO. The nuclei were counterstained with Hoechst33342. The luminescence of all color images was analyzed as described above. Data are the mean ± SD (n=8). Data were analyzed by own-way analysis of variance followed by Tuckey’s post hoc test.\*\*\**P* <0.001 vs. control. ###*<u>P</u>* <0.001. Bar = 300 μm. (D) HOC-313 cells were pretreated with catalase for 1h and then incubated with ODM for 2h. Mitochondria, intracellular hydroxyl radicals, and nuclei were stained with MTG, OXO, and Hoechst33342. Bar =20 μm.

### 3.5. Reduced H_2_O_2_ removal augments ODM-induced H_2_O_2_ increase and cell death in tolerant cells

We noticed that the effect of ODM decreased as cell density increased. As a result, HOC-313 cells grown at a high density became highly resistant to treatment with ODM (50%) (Fig. 5A). As the above results suggested that H_2_O_2_ was critical in oxidative cell death, we assumed the resistance might be related to H_2_O_2_-scavenging activity. To test this hypothesis, we examined the impact of agents affecting the activity on ODM cytotoxicity. The glutathione (GSH) synthase inhibitor BSO has been shown to reduce GSH synthesis and GSH-dependent antioxidant systems, including GSH peroxidase (GPX), the primary H_2_O_2_-scavenging enzyme [30, 31]. On the other hand, 3-AT, an irreversible catalase inhibitor, can block H_2_O_2_ removal by catalase. BSO had moderate cytotoxicity and markedly augmented growth inhibition and cell death caused by ODM (Fig. 5A, B). BSO also synergistically increased intracellular H_2_O_2_ with ODM (Fig. 5C). 3-AT also augmented the growth inhibitory effect to a lesser extent. These results suggest that the resistance to ODM may be due to increased H_2_O_2_ removal.

**Figure 5.**
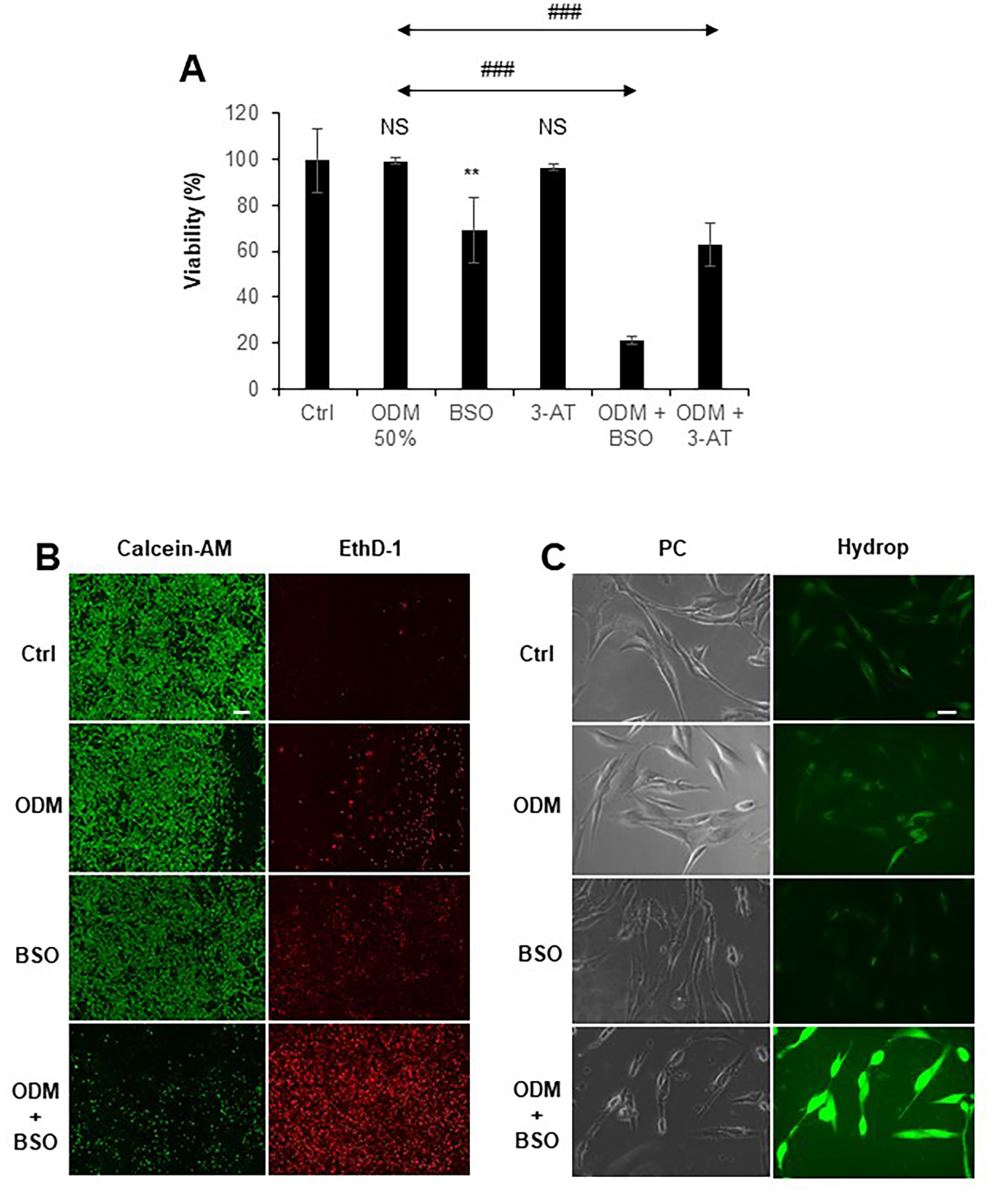
BSO augments ODM-induced H_2_O_2_ increase and cell death in insensitive cells. (A) HOC-313 cells were pretreated with the DL-Buthionine-(S, R)-sulfoximine (BSO, 100 μM) or 3-AT (1 mM) alone or in combination for 1h. then they were treated with the ODM (50%) for 72h. Cells were analyzed for viability as described in the legend of Figure 3. Data were analyzed by own-way analysis of variance followed by Tuckey’s post hoc test. Data are the mean ± SD (n=7 or 8). \*\**P* < 0.01; NS, not significant vs. control; ###*P* <0.001. (B) The cells were pretreated with BSO for 1h and then treated with ODM (50%). They were incubated for 18h and analyzed for cell death using Calcein-AM (Calcein, 2 μM) and ethidium bromide-1 (EthD-1, 2 μM). Live cells were stained green with Calcein, whereas dead/dying cells were stained red with EthD-1. Images were obtained from BZ-X 710 Digital Biological Microscope with a 4x objective. Bar = 300 μm. (C) The cells were treated with the ODM (50%) or BSO alone or in combination for 18h and were stained with HYDROP™ (Hydrop,1μM). Images were obtained from BZ-X 710 Digital Biological Microscope with a 40x objective and analyzed as previously described in the legend of Figure 4. Bar =20 μm.

### 3.6. ODM induces MPMC and microtubule remodeling in an H_2_O_2_-dependent manner

In untreated cells, most mitochondria belong to Type I or II. Following ODM treatment, they became Type III (Fig. 6A). These morphological changes occurred as rapidly as within 2 h after ODM treatment. Like ODM cytotoxicity, catalase prevented the effect. Quantitative analyses of Type I, II, and III mitochondria ratios confirmed these observations (Fig. 6B). Similar morphological changes were observed in SAS, HOS, and U251MG (Supplementary Fig. S1A–C). The above results suggested the onset of mitochondrial movement before the distribution change. Since mitochondria can move along the microtubule track, we analyzed the impact of ODM on tubulin dynamics. In untreated cells, tubulin had a network structure distributing broadly on both sides of the nuclei in the cytoplasm (Pan-cytoplasmic) (Fig. 6C). On the other hand, after ODM treatment, it condensed and assembled one side of the nuclei (Perinuclear). Catalase also prevented the effect (Fig. 6C). Similarly, the microtubule inhibitor Nocodazole (NC) prevented MPMC and the distribution shift (Supplementary Fig. S1D). These results show that ODM induces MPMC and microtubule remodeling in an H_2_O_2_-dependent manner.

**Figure 6.**
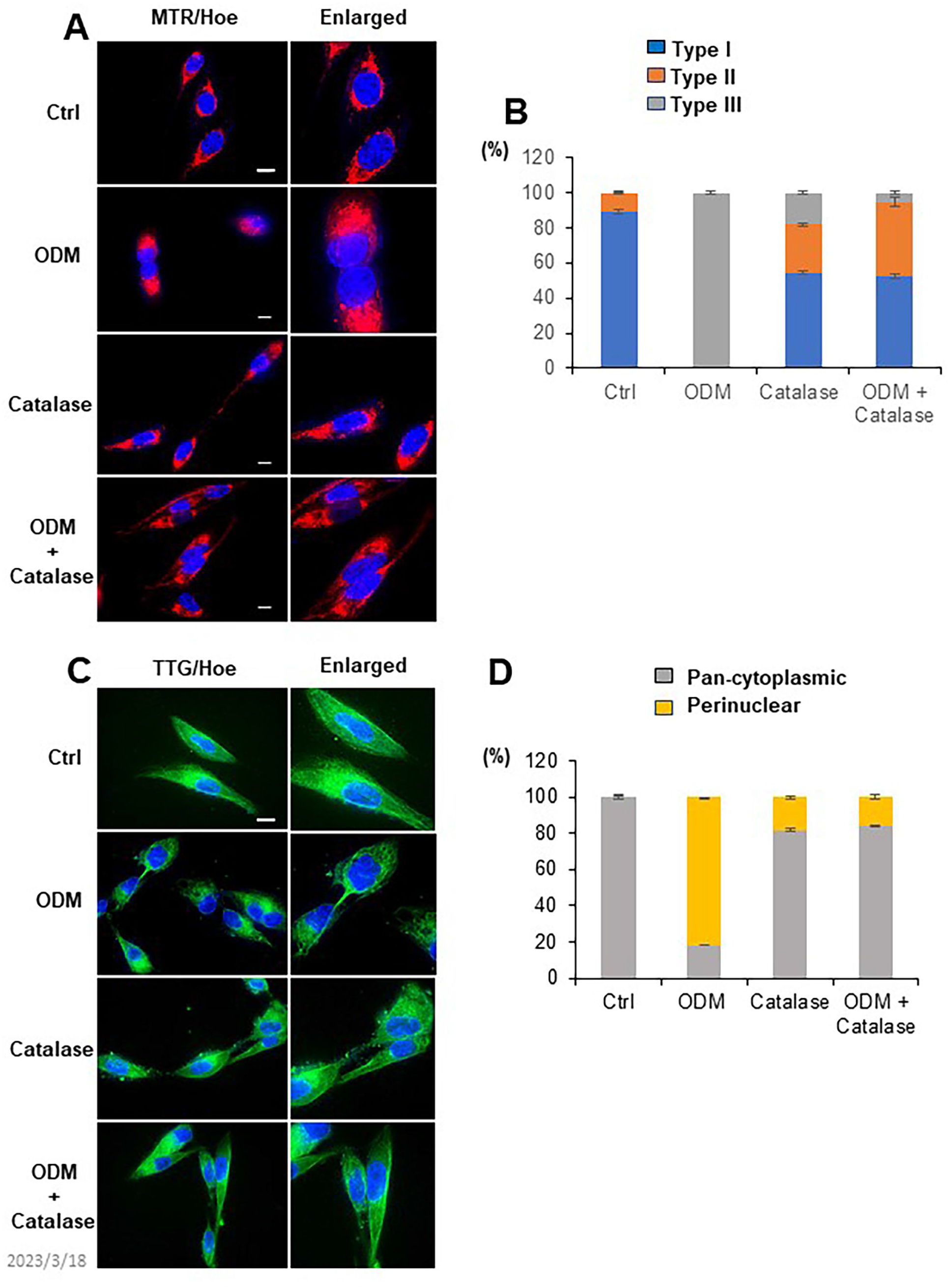
ODM induces MPMC and tubulin remodeling in an H_2_O_2_-dependent manner. (A) After treating HOC-313 cells with medium or ODM (50 %) for 2h, mitochondria and nuclei were stained with 20 nM MTR and Hoechest33342, respectively. Images were obtained from BZ-X 710 Digital Biological Microscope with a 100x objective and analyzed using BZ-H3A application software. Bar =10 μm. (B) Mitochondria exhibiting three different subcellular distributions, Pan-cytoplasmic (Type I), PNMC (Type II), and MPMC (Type III), were counted in two or three pictures. Data are the mean± SD (n=∼20). (C) The cells were treated with ODM (50%) for 2h and stained with Tubulin Tracker™ Green (TTG, 100 nM) for 30 min. Bar =10 μm. (D) Tubulin exhibiting Pan-cytoplasmic or Perinuclear distribution was counted in two or three pictures. Data are the mean± SD (n=∼20).

### 3.7. ODM causes minimal MPMC, ROS production, and cell death in non-malignant cells

Next, we examined whether ODM had cytotoxicity in non-malignant cells. ODM (≤25%) had minimal effect on the viability of HaCaT and hFOB, the non-transformed counterparts of OC and OS, respectively (Fig. 7A, B). Similar results were obtained in HDFs (not shown). At a higher concentration (50%), ODM decreased cell viability moderately (around 50%). Moreover, minimal cell and nuclear morphological changes were observed after ODM treatment in HaCaT and HDFs (Fig. 7C, D). ODM had a minimal effect on mitochondrial morphology in HaCaT while increasing mitochondrial fragmentation but evoked MPMC minimally in HDFs (Fig. 7C, D). In addition, ODM minimally increased intracellular H_2_O_2_ and ^·^OH in HaCaT cells (Supplementary Fig. S2). These results indicate that ODM causes minimal MPMC, ROS production, and cell death in non-malignant cells.

**Figure 7.**
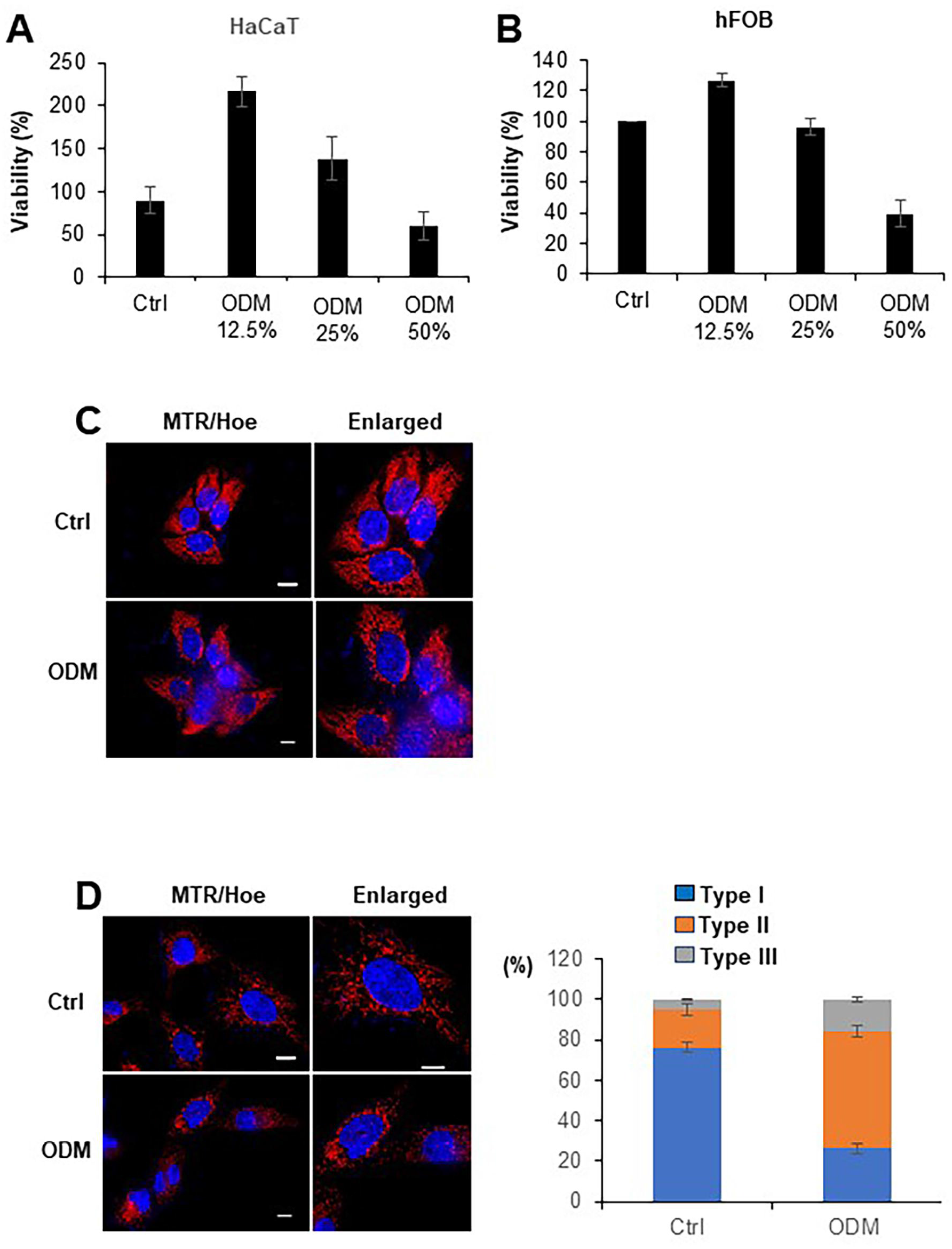
ODM has low cytotoxicity in non-malignant cells. (A, B) HaCaT (A) and hFOB (C) cells were treated with ODM at the indicated concentration for 72h and were measured for viability by a WST-8 assay. Data were analyzed by own-way analysis of variance followed by Tuckey’s post hoc test. \*\**P* < 0.01; NS, not significant vs. control. (C, D) HaCaT (C) and HDFs (D) were treated with ODM (50%) for 2h. After stimulation, cells were stained with MTR and Hoechst33342. Images were obtained and analyzed as described in the legend of Figure 6. Bar =10 μm. Mitochondria exhibiting three different subcellular distributions (Type I, Type II, Type III) were counted in two or three pictures. Data are the mean± SD (n=∼20).

## 4. Discussion

The present study aimed to examine the antitumor activity of dO_3_ and verify its role in mediating the action of APAM. As expected, ODM had considerable amounts of O_3_ (Fig. 1) and potent cytotoxicity in various cancer cells, including OC, OS, and GBM (Fig. 2). Moreover, it primarily triggered oxidative cell death in OC cells. Meanwhile, apoptosis explicitly participated in the moderate effect (Fig. 3). Thus, apoptosis induction may occur under limited conditions and may be insufficient for killing OC cells effectively. Oxidative cell death may be essential for efficient OC killing. In line with the role of oxidative stress, ODM increased mitochondrial ROS, such as O_2_^·-^ and ^·^OH (Fig. 4). catalase prevented the increases while reduced H_2_O_2_-removal upregulated them (Fig. 4 and 5). These findings indicate that H_2_O_2_ mediates oxidative stress. These results are similar to those obtained with APAM previously described [28]. Moreover, APAM had dO_3_ as much as ODM (Fig. 1). APAM has potent antitumor activity against different cancer cell lines. More importantly, it acts on cancer cells preferentially [28]. Notably, our data indicate that ODM has similar tumor-selective action. ODM showed little cytotoxicity against non-malignant cells and minimally increased MPMC and ROS production (Fig. 7). Our results indicate that dO_3_ has tumor-selective cytotoxicity and support the view that dO_3_ mediates the action of APAM. In addition, ODM had NO_2_^-^ and NO_3_^-^ below the detection limit (Fig. 1), indicating that the effect is independent of NOx.

It is known that dO_3_ can directly generate H_2_O_2_ in the reaction with H_2_O. Our previous study demonstrated that H_2_O_2_ increased mitochondrial O_2_^·-^ via the reduction of the electron transport chain in different cancer cells [32]. In addition, H_2_O_2_ also can generate ^·^OH in reacting with iron (II) or copper (I). Collectively, H_2_O_2_ produced from dO_3_ can trigger mitochondrial oxidative stress. Another finding that GSH synthesis inhibition sensitized tolerant cells to ODM (Fig. 5) supports this view. These observations suggest that the GSH-dependent H_2_O_2_-removal mechanism, possibly GPXs, is vital in removing H_2_O_2_ after dO_3_ treatment. The inhibition of catalase activity by AT increased mitochondrial ROS but minimally affected cell survival (Fig. 4 and 5), suggesting the requirement of another event to cause cell death in tolerant cells. Ferroptosis has emerged as a critical oxidative cell death mode in cancer cells. It is regulated necrosis resulting from LPO accumulation where GPX and iron are crucial (for a recent review, see [33]). ODM could increase ^·^OH, the initiator of lipid peroxidation (Fig. 4). In addition, preliminary results showed that iron chelation partially prevented the effect, while iron (II) addition augmented it. Therefore, ^·^OH generation from H_2_O_2_ and iron (II) via the Fenton reaction might play a role in the production. This view favors the notion that besides H_2_O_2_, other factors are essential for sufficient cell death. Our data indicate similarities and differences between cell death caused by ODM and ferroptosis. The possible involvement of GPX4 resembles ferroptosis. On the other hand, the failure of Fer-1 in protecting cells from ODM (Fig. 3) suggests the role of different mechanisms and lipid radicals. Further characterization, including the role of several master ferroptosis regulators and genes, is underway.

Our results showed that similar to APAM [28], ODM could cause MPMC and tubulin remodeling (Fig. 6). The findings provide evidence for the role of dO_3_ in the action of APAM. Notably, catalase blocked these two events, indicating the critical role of H_2_O_2_ in regulating MPMC and tubulin remodeling. MPMC may involve multiple sequential cellular events, mitochondrial fragmentation, movement, and assembly. In line with the role of H_2_O_2_, we previously demonstrated that H_2_O_2_ could cause mitochondrial fragmentation in MM cells via increased mitochondrial O_2_^·-^ [34] (Saito et al., 2016). This event was associated with increased phosphorylation of Drp1 at Ser 616, the driving signal of mitochondrial fission. Similar mechanisms might be involved in the effect of ODM. Tubulin polymerization is critical in altered mitochondrial distribution from Type I to Type II in response to stresses, such as hypoxia and heat shock [26, 27]. This event is required for the mitochondrial movement along the microtubule track by motor proteins such as Kinesin and Dynein. Our data indicate that catalase (Fig. 6) and NC (Supplementary Fig. S1D) can prevent tubulin remodeling and MPMC caused by ODM, suggesting the involvement of H_2_O_2_ and tubulin polymerization in the mitochondrial movement. Besides mitochondrial oxidative stress, plasma membrane depolarization (PMD) is essential for mitochondrial assembly [35]. Notably, dO_3_ can affect biological systems in several different forms. It can react with target biomaterials directly and specifically as O_3_ itself. It can also attack poly unsatisfied fatty acids (PUFAs) in the cell membrane, resulting in lipid oxides. The production follows PUFA peroxidation and the production of other toxic radicals and substances, such as LO^·^, LOO^·^, lipo hydroperoxides (LOOH), hydroxy-2,3 trans-noneal (HNE), and malonyl dialdehyde (MDA) [13]. These radicals and aldehydes are toxic and could contribute to ODM cytotoxicity. Notably, linoleic acid hydroperoxide can evoke the depolarization of plasma membrane and mitochondrial membrane potentials [36]. Therefore, LPOs like lipid hydroperoxides might trigger PMD and mitochondrial assembly. Further studies to explore this possibility are ongoing.

Several prior studies have implicated the role of O_3_ in the antitumor effect of PTLs. Mokhtari and colleagues [37] have shown the production of O_3_ in plasma-activated media and the correlation between the amount and antitumor activity. However, this report lacks evidence for the oxidant’s role in the media’s action. Lunov et al. [38, 39] have demonstrated that O_3_ is an abundant component of air plasma and that O_3_ gas flush induces necrosis. Their findings are similar to the present findings, while there are some discrepancies between our results and theirs. The authors reported the highest toxicity of O_3_ gas for non-malignant cells. In contrast, dO_3_ had little toxicity in the present study (Fig. 7). The report also demonstrated that O_3_ gas caused mitochondrial permeability transition. However, our preliminary results showed that dO_3_ had no such effect. A possible explanation for the discrepancies may be the different biological effects between O_3_ gas and dO_3_. As described above, dO_3_ can exhibit its physical impacts by acting in various forms other than O_3_.

In summary, this study shows that dO_3_ has tumor-selective cytotoxicity primarily via oxidative cell death independently of NOx. Our results suggest that mitochondrial ROS and the resulting tubulin remodeling and MPMC play a vital role in the action. Also, our data support the view that dO_3_ is a critical mediator of the action of APAM. Thus, ODM could be a more chemically-defined alternative to PTLs in cancer treatment.

## Abbreviations

7-AAD: 7-amino-actinomycin D
APAM: air plasma-activated medium
CAP: cold atmospheric plasma
CL: cardiolipin
Drp: dynamin-related protein
Fer-1: ferrostatin-1
Hoe,mn: Hoechst 33342
LPO: lipid peroxide
MitoSOX: MitoSOX Red CMXRos
MPMC: monopolar perinuclear mitochondrial clustering
mROS: mitochondrial reactive oxygen species
MTR: MitoTracker Red
nROS: nuclear reactive oxygen species
NAC: N-acetylcysteine
NAO: 10-N-nonyl acridine orange
NAC: N-acetylcysteine
NC: Nocodazole
OC: oral cancer
OS: osteosarcoma
PNMC: perinuclear mitochondrial clustering
TTG: tubulin tracker green.

## Supplementary Figure legends

**Figure S1. ODM induces MPMC and tubulin remodeling in different tumor cells via microtubules**. (A–C) SAS (A), HOS (B), and U251MG cells (C) were treated with medium or ODM (50 %) for 2h, and mitochondria and nuclei were stained with 20 nM MTR and Hoechest33342, respectively. Images were obtained from BZ-X 710 Digital Biological Microscope with a 100x objective and analyzed. Bar =10 μm. (D) HOC-313 cells were treated with ODM (50%) for 2h and stained with MTR, TTG, and Hoechest33342. Images were taken and analyzed as described in the legend of Figure 6. Bar =10 μm.

**Figure S2. ODM minimally increases intracellular ROS in non-malignant cells**. HaCaT cells were treated with ODM (50%) for 2h. After stimulation, H_2_O_2_ and hydroxyl radicals were stained with HYDROP™ (Hydrop,1μM) and OxiORANGE (OXO, 1 μM), respectively. The nuclei were stained with Hoechst33342. Images were obtained from BZ-X 710 Digital Biological Microscope with a 40x objective and analyzed as described above. Bar = 20 μ_m_.

## Acknowledgments

We thank the JCRB Cell Bank of the National Institutes of Biomedical Innovation, Health, and Nutrition (Osaka, Japan) and the Riken BioResource Center (Tsukuba, Japan) for providing cell lines.

## CrediT authorship contribution statement

Conceptualization, Y. S.-K., M. S.-K. (Manami Suzuki-Karasaki), S.I. H.O. investigation, M. S.-K. (Manami Suzuki-Karasaki), M. S.-K. (Miki Suzuki-Karasaki), Y.O., S.I., H.O. Data acquisition and analysis, M. S.-K. (Manami Suzuki-Karasaki), M. S.-K. (Miki Suzuki-Karasaki), Y.O., S.I., H.O. Funding acquisition, Y. S.-K., M. S.-K. (Manami Suzuki-Karasaki). Methodology, S.I., H.O. Visualization, Y. S.-K., M. S.-K. (Manami Suzuki-Karasaki). Project administration, Y. S.-K., S.I., H.O. Resources, S.I., H.O., H.N. writing–original draft preparation, Y. S.-K., M. S.-K. (Manami Suzuki-Karasaki). writing–review and editing, Y. S.-K., M. S.-K. (Manami Suzuki-Karasaki), H.N. supervision, Y. S.-K., H.N.

## References

1. Karunakaran K, Muniyan R. Genetic alterations and clinical dimensions of oral cancer: a review. Mol Biol Rep. 2020;47(11):9135–48.

2. Sha J, Bai Y, Ngo HX, Okui T, Kanno T. Overview of Evidence-Based Chemotherapy for Oral Cancer: Focus on Drug Resistance Related to the Epithelial-Mesenchymal Transition. Biomolecules. 2021;11(6).

3. Kornienko A, Mathieu V, Rastogi SK, Lefranc F, Kiss R. Therapeutic agents triggering nonapoptotic cancer cell death. J Med Chem. 2013;56(12):4823–39.

4. Keidar M, Walk R, Shashurin A, Srinivasan P, Sandler A, Dasgupta S, et al. Cold plasma selectivity and the possibility of a paradigm shift in cancer therapy. Br J Cancer.

5. Yan D, Sherman JH, Keidar M. Cold atmospheric plasma, a novel promising anticancer treatment modality. Oncotarget. 2017;8(9):15977–95.

6. Ando T, Suzuki-Karasaki M, Ichikawa J, Ochiai T, Yoshida Y, Haro H, et al. Combined Anticancer Effect of Plasma-Activated Infusion and Salinomycin by Targeting Autophagy and Mitochondrial Morphology. Front Oncol. 2021;11:593127.

7. Ito T, Ando T, Suzuki-Karasaki M, Tokunaga T, Yoshida Y, Ochiai T, et al. Cold PSM, but not TRAIL, triggers autophagic cell death: A therapeutic advantage of PSM over TRAIL. Int J Oncol. 2018;53(2):503–14.

8. Yoshikawa N, Liu W, Nakamura K, Yoshida K, Ikeda Y, Tanaka H, et al. Plasma-activated medium promotes autophagic cell death along with alteration of the mTOR pathway. Sci Rep. 2020;10(1):1614.

9. Jo A, Bae JH, Yoon YJ, Chung TH, Lee EW, Kim YH, et al. Plasma-activated medium induces ferroptosis by depleting FSP1 in human lung cancer cells. Cell Death Dis. 2022;13(3):212.

10. Girard PM, Arbabian A, Fleury M, Bauville G, Puech V, Dutreix M, et al. Synergistic Effect of H2O2 and NO2 in Cell Death Induced by Cold Atmospheric He Plasma. Sci Rep. 2016;6:29098.

11. Bauer G. The synergistic effect between hydrogen peroxide and nitrite, two long-lived molecular species from cold atmospheric plasma, triggers tumor cells to induce their cell death. Redox Biol. 2019;26:101291.

12. Baeza-Noci J, Pinto-Bonilla R. Systemic Review: Ozone: A Potential New Chemotherapy. Int J Mol Sci. 2021;22(21).

13. Bocci VA. Scientific and medical aspects of ozone therapy. State of the art. Arch Med Res. 2006;37(4):425–35.

14. Bocci V, Travagli V, Zanardi I. Randomised, double-blinded, placebo-controlled, clinical trial of ozone therapy as treatment of sudden sensorineural hearing loss. J Laryngol Otol. 2009;123(7):820; author reply

15. Clavo B, Santana-Rodríguez N, Llontop P, Gutiérrez D, Suárez G, López L, et al. Ozone Therapy as Adjuvant for Cancer Treatment: Is Further Research Warranted? Evid Based Complement Alternat Med. 2018;2018:7931849.

16. Zänker KS, Kroczek R. In vitro synergistic activity of 5-fluorouracil with low-dose ozone against a chemoresistant tumor cell line and fresh human tumor cells. Chemotherapy. 1990;36(2):147–54.

17. Cannizzaro A, Verga Falzacappa CV, Martinelli M, Misiti S, Brunetti E, Bucci B. O(2/3) exposure inhibits cell progression affecting cyclin B1/cdk1 activity in SK-N-SH while induces apoptosis in SK-N-DZ neuroblastoma cells. J Cell Physiol. 2007;213(1):115–25.

18. Simonetti V, Quagliariello V, Giustetto P, Franzini M, Iaffaioli RV. Association of Ozone with Fluorouracil and Cisplatin in Regulation of Human Colon Cancer Cell Viability: In Vitro Anti-Inflammatory Properties of Ozone in Colon Cancer Cells Exposed to Lipopolysaccharides. Evid Based Complement Alternat Med. 2017;2017:7414083.

19. Kamm A, Przychodzen P, Kuban-Jankowska A, Jacewicz D, Dabrowska AM, Nussberger S, et al. Nitric oxide and its derivatives in the cancer battlefield. Nitric Oxide. 2019;93:102–14.

20. Kim J, Thomas SN. Opportunities for Nitric Oxide in Potentiating Cancer Immunotherapy. Pharmacol Rev. 2022;74(4):1146–75.

21. Twig G, Shirihai OS. The interplay between mitochondrial dynamics and mitophagy. Antioxid Redox Signal. 2011;14(10):1939–51.

22. Hoppins S, Nunnari J. Cell Biology. Mitochondrial dynamics and apoptosis--the ER connection. Science. 2012;337(6098):1052–4.

23. Pendin D, Filadi R, Pizzo P. The Concerted Action of Mitochondrial Dynamics and Positioning: New Characters in Cancer Onset and Progression. Front Oncol. 2017;7:102.

24. Pedriali G, Rimessi A, Sbano L, Giorgi C, Wieckowski MR, Previati M, et al. Regulation of Endoplasmic Reticulum-Mitochondria Ca. Front Oncol. 2017;7:180.

25. Quintana A, Schwarz EC, Schwindling C, Lipp P, Kaestner L, Hoth M. Sustained activity of calcium release-activated calcium channels requires translocation of mitochondria to the plasma membrane. J Biol Chem. 2006;281(52):40302–9.

26. Al-Mehdi AB, Pastukh VM, Swiger BM, Reed DJ, Patel MR, Bardwell GC, et al. Perinuclear mitochondrial clustering creates an oxidant-rich nuclear domain required for hypoxia-induced transcription. Sci Signal. 2012;5(231):ra47.

27. Agarwal S, Ganesh S. Perinuclear mitochondrial clustering, increased ROS levels, and HIF1 are required for the activation of HSF1 by heat stress. J Cell Sci. 2020;133(13).

28. Suzuki-Karasaki M, Ando T, Ochiai Y, Kawahara K, Nakayama H, Suzuki-Karasaki Y. Air Plasma-Activated Medium Evokes a Death-Associated Perinuclear Mitochondrial Clustering. Int J Mol Sci. 2022;23(3).

29. Suzuki Y, Inoue T, Murai M, Suzuki-Karasaki M, Ochiai T, Ra C. Depolarization potentiates TRAIL-induced apoptosis in human melanoma cells: role for ATP-sensitive K+ channels and endoplasmic reticulum stress. Int J Oncol. 2012;41(2):465–75.

30. Meister A. Mitochondrial changes associated with glutathione deficiency. Biochim Biophys Acta. 1995;1271(1):35–42.

31. Jiang H, Wang H, De Ridder M. Targeting antioxidant enzymes as a radiosensitizing strategy. Cancer Lett. 2018;438:154–64.

32. Inoue T, Suzuki-Karasaki Y. Mitochondrial superoxide mediates mitochondrial and endoplasmic reticulum dysfunctions in TRAIL-induced apoptosis in Jurkat cells. Free Radic Biol Med. 2013;61:273–84.

33. Stockwell BR. Ferroptosis turns 10: Emerging mechanisms, physiological functions, and therapeutic applications. Cell. 2022;185(14):2401-21.34.

34. Saito K, Asai T, Fujiwara K, Sahara J, Koguchi H, Fukuda N, et al. Tumor-selective mitochondrial network collapse induced by atmospheric gas plasma-activated medium. Oncotarget. 2016;7(15):19910–27.

35. Suzuki-Karasaki Y, Fujiwara K, Saito K, Suzuki-Karasaki M, Ochiai T, Soma M. Distinct effects of TRAIL on the mitochondrial network in human cancer cells and normal cells: role of plasma membrane depolarization. Oncotarget. 2015;6(25):21572–88.

36. Forman HJ, Kim E. Inhibition by linoleic acid hydroperoxide of alveolar macrophage superoxide production: effects upon mitochondrial and plasma membrane potentials. Arch Biochem Biophys. 1989;274(2):443–52.

37. Mokhtari H, Farahmand L, Yaserian K, Jalili N, Majidzadeh-A K. The antiproliferative effects of cold atmospheric plasma-activated media on different cancer cell lines, the implication of ozone as a possible underlying mechanism. J Cell Physiol. 2019;234(5):6778–82.

38. Lunov O, Zablotskii V, Churpita O, Chánová E, Syková E, Dejneka A, et al. Cell death induced by ozone and various non-thermal plasmas: therapeutic perspectives and limitations. Sci Rep. 2014;4:7129.

39. Lunov O, Zablotskii V, Churpita O, Lunova M, Jirsa M, Dejneka A, et al. Chemically different non-thermal plasmas target distinct cell death pathways. Sci Rep. 2017;7(1):600.

