## Supplementary figures and images for "Ozone mediates tumor-selective cell death caused by air plasma-activated medium independently of NOx"

### S1

Supplementary
Fig. S1

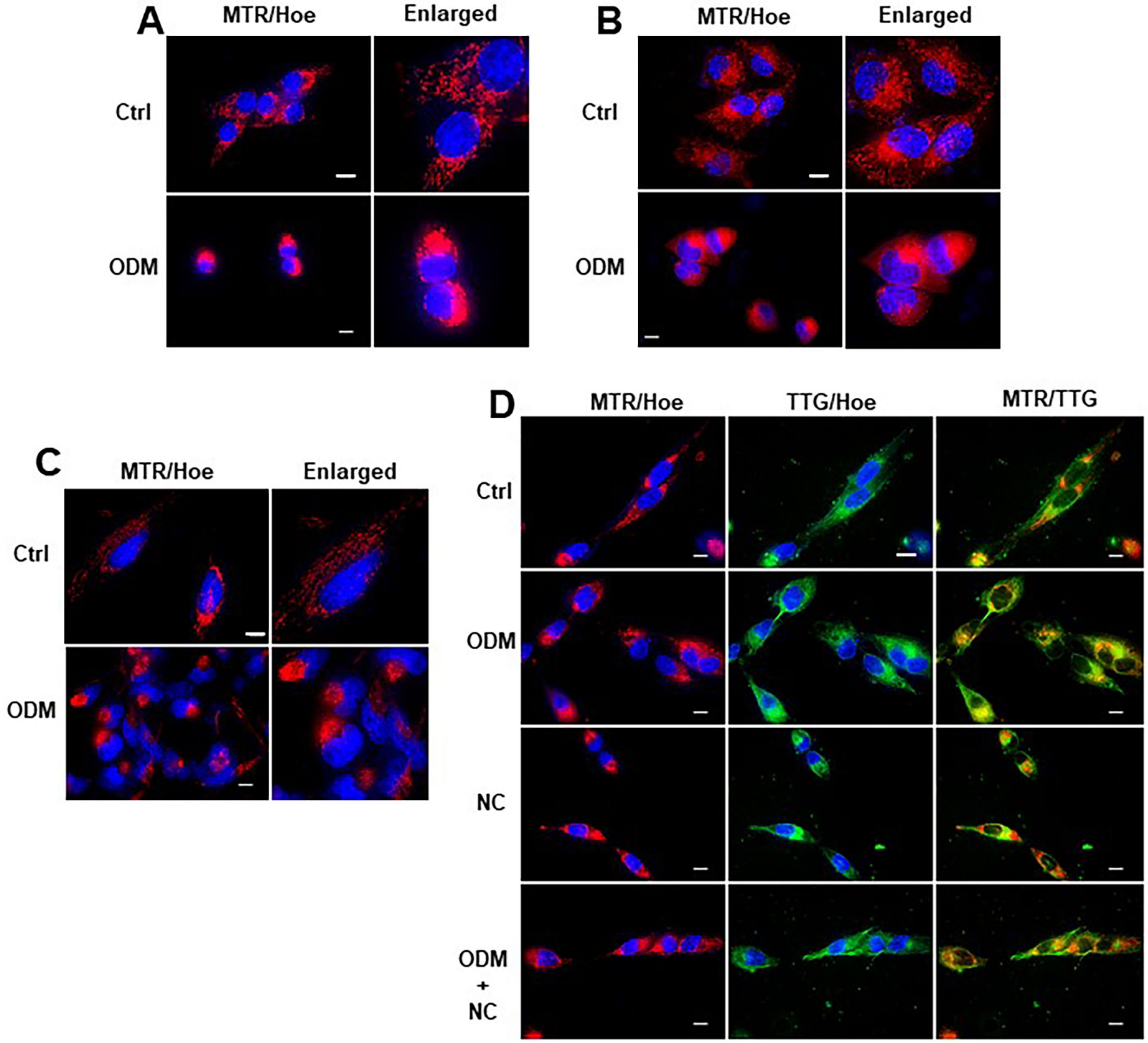


2023/3/18

### S2

Supplementary
Fig. S2

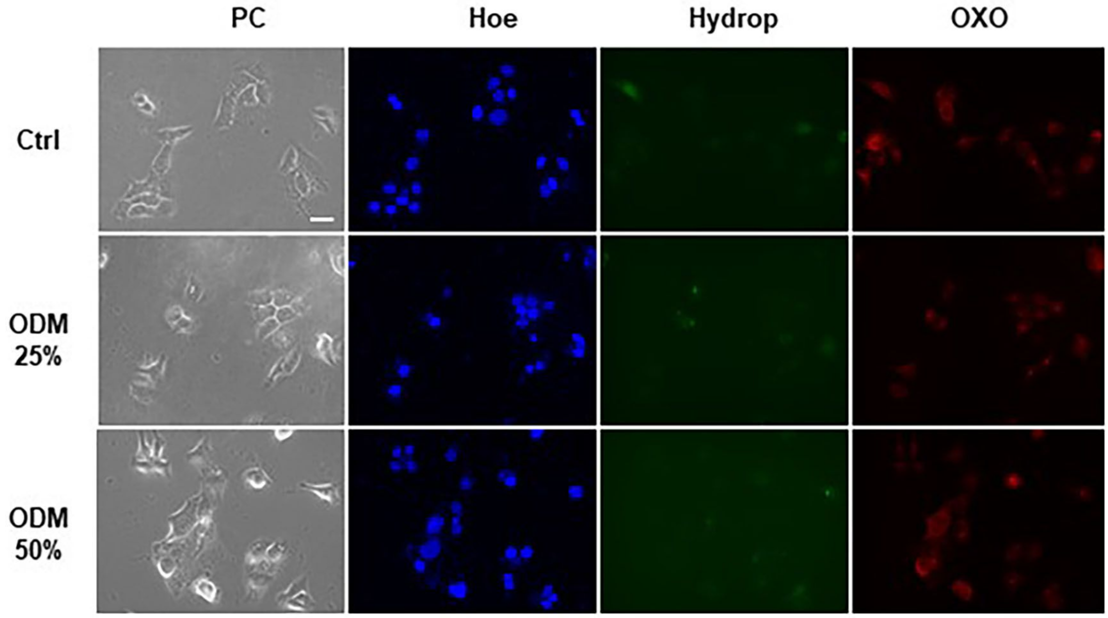
